# Amplification of disease by nutrient addition: Testing mechanisms from individual to community levels

**DOI:** 10.1101/2024.10.08.617235

**Authors:** Elizabeth T. Green, Robert W. Heckman, Charles E. Mitchell

## Abstract

Nutrient supply can amplify disease epidemics through mechanisms from individual to community levels. Within host individuals, nutrient addition can drive pathogen replication or growth. Across a host population, nutrient addition can drive disease transmission by increasing host growth and abundance relative to defense. Furthermore, such effects may be influenced by pathogen species interactions. Understanding how nutrients impact disease epidemics requires a framework that integrates these mechanisms across biological levels. To build such a framework, we conducted a field experiment in an old field on tall fescue, *Lolium arundinaceum*, and used structural equation models to integrate multiple hypothesized mechanisms. Nutrient addition (NPK fertilizer) increased brown patch disease but was best modeled as a direct path and not mediated by host abundance. To expand our framework, we also re-analyzed a previous experiment. That experiment reproduced the direct path from nutrients to disease, and added an indirect path mediated by host population abundance. Nutrient addition also increased foliar nitrogen, consistent with individual-level mechanism, but this did not increase disease. Brown patch decreased with burden of another disease, anthracnose, independently of nutrients. These results partially support both individual- and population-level hypotheses, emphasizing the importance of considering multiple biological levels underlying impacts of abiotic change.

## Introduction

Nutrients play an important role in structuring ecological communities (Borer et al. 2014, Arnillas et al. 2021) and understanding this role is of rising importance as human activities increase the supplies of nutrients to many communities. The availability of nutrients to plant hosts may impact interactions between plants and their diverse herbivores and pathogens by two main hypothesized mechanisms, the growth-defense tradeoff hypothesis and the nutrient-supply hypothesis (Mitchell et al. 2003, Zaret et al. 2024). These two hypotheses are not mutually exclusive; they predict the effects of nutrient addition on disease at different levels of organization. Individual plants, the subject of the nutrient-supply hypothesis, are embedded within a greater community of potential hosts, which is the focus of the growth-defense trade-off hypothesis. Transmission of pathogens can be modulated by processes at both the scales of the individual host and of the host community. Incorporating theory and study of both levels of organization, allows for a deeper understanding of the pathogen infection and spread (Borer et al. 2016, Miller et al. 2018, Bolnick et al. 2020).

The growth-defense tradeoff hypothesis states that plants facing infection can allocate resources to either growth or mounting a defense against disease (Lind et al. 2013, Züst and Agrawal 2017, Zaret et al. 2024). Fast growing plants do so at the expense of defense, leaving them vulnerable to disease (Liu et al. 2017, Heckman et al. 2019). A community dominated by fast growing, poorly defended plants may then experience an increase in disease burden and damage. At the organizational scale of the plant community, nutrient addition can cause a shift from slow to fast growing species, which can lead to an increase in aboveground plant biomass (Wright and Fridley 2010, Halliday et al. 2020a, Heckman 2023). The growth-defense hypothesis predicts that these fast growing plants will experience an increased disease burden (Cappelli et al. 2020, Zaret et al. 2024). Additionally, increasing abundance of poorly defended plants can increase disease transmission through a community (Mordecai 2011, Halliday et al. 2021).

Individual plants also have phenotypic plasticity in response to nutrient supply. Plants can alter plant tissue concentrations, biochemical defenses, and growth under different nutrient availabilities (Zandt 2007). The nutrient-disease hypothesis proposes that plants growing under heightened nutrient availability may experience increased disease due to increased leaf nitrogen availability to pathogens (Dordos 2008, Veresoglou et al. 2013, Ebeling et al. 2021). At the scale of the individual host, nutrient addition can increase the disease burden directly through increasing pathogen growth due to increased host quality (Veresoglou et al. 2013, Heckman et al. 2019).

In both wild and agricultural systems, plant populations and individuals often face simultaneous infection pressure from multiple pathogens (Abdullah et al. 2017, Halliday et al. 2020a, O’Keeffe et al. 2021a, Grunberg et al. 2023b). Ecological interactions among pathogen species can range from competition to facilitation, with the potential to dampen or amplify disease epidemics (Halliday et al. 2020b, O’Keeffe et al. 2021b). Thus, pathogen species interactions create the potential for additional indirect effects of nutrient supply on disease epidemics. Yet, few experimental tests of either the nutrient-disease hypothesis or the growth-defense hypothesis have explicitly considered ecological interactions between pathogen species (Marchetto and Power 2018, Halliday et al. 2019, Zaret et al. 2024).

To address this gap, we conducted a multi-year field experiment and re-analyzed data from a previous experiment at the same site, applying structural equation modeling to evaluate the links between soil nutrients, the plant community, and three diseases as well as herbivory on the community’s dominant plant species, tall fescue, *Lolium arundinaceum*. Previous studies have found tall fescue to be highly nutrient responsive and to experience high rates of herbivory and disease in the field relative to co-occurring species, part of a community-level tradeoff between growth and defense (Heckman et al. 2019). Based on the growth-defense hypothesis, we predicted that nutrient addition would increase total plant biomass and abundance of tall fescue, a fast-growing host plant, and thereby indirectly increase tall fescue’s disease burden. In keeping with the nutrient-disease hypothesis, we predicted that nutrient addition would increase leaf nitrogen concentration of tall fescue and thereby indirectly increase its disease burden. We further predicted that increased disease burden of anthracnose would facilitate brown patch, as shown in previous lab inoculation experiments with tall fescue (O’Keeffe et al. 2021b, Green et al. 2024), creating an additional indirect positive effect of nutrient addition on brown patch. By testing these individual-level and community-level hypotheses in an integrated framework, this study aimed to help unify our understanding of nutrient effects on disease in ecological communities.

## Methods

### Study System

Tall fescue (*Lolium arundinaceum*) is a dominant grass species in non-cultivated systems across the southern and eastern United States and is host to numerous fungal pathogens (Halliday et al. 2019, 2020a). In this experiment, we focus on three foliar fungal pathogens that frequently coinfect their host and differ substantially in biology. First, *Rhizoctonia solani* exhibits a necrotrophic strategy of killing leaf tissue and extracting nutrients from the dead or dying tissue. It disperses via soil and water splash and causes the disease brown patch (Ogoshi 1987). Second, *Colletotrichum cereale* is a hemibiotrophic parasite, initially infecting and extracting nutrients from living tissue, then killing leaf tissue and causing necrotic lesions. It disperses via water droplets and causes the disease anthracnose (Crouch and Beirn 2009). Finally, *Puccinia coronata* utilizes a biotrophic strategy of infecting and extracting nutrients from living tissue. It disperses via wind-borne spores and causes the disease crown rust (Liu and Hambleton 2013).

### Field setup

The experiment was conducted across three growing seasons at Widener Farm (Duke Forest Teaching and Research Laboratory), an 8-ha old field located in Orange County, North Carolina that was last cultivated in 1996 (Heckman et al. 2016). The site is dominated by perennial grasses, including tall fescue (*Lolium arundinaceum*). We staked out 72 1m × 1m field plots in a 6 × 12 plot array, then sprayed every other plot with glyphosate herbicide (Roundup® 360, Bayer) on May 2, 2021, and covered the sprayed plots with landscaping fabric to kill all aboveground vegetation the following week. This produced a checkerboard pattern of 36 sprayed plots that lacked standing vegetation. We separated the 36 sprayed plots into three spatial blocks of 12 plots each and randomly assigned each plot to one of three treatments of 10-10-10 N-P-K Meherrin Fertilizer (no fertilizer, low fertilizer (5g/m^2^), high fertilizer (10g/m^2^)). Each fertilizer treatment was replicated four times in each block. The plots were treated with their assigned nutrient addition in the spring of each year.

We transplanted tall fescue plants into these field plots and allowed for natural recolonization of other species from the seedbank and surrounding vegetation. To do this, early April of 2021, we primed and germinated tall fescue seed collected from Widener Farm in 2018. Seed was primed by soaking in water for six hours and then left out to dry overnight. We then sprinkled the seed onto moist vermiculite and left it in a closed Tupperware container for 10 days. After that, we transferred the seeds to individual pots filled with MetroMix® 360 soil (SunGro). In the greenhouse, we watered seedlings three times per week for eight weeks, at which point plants were hardy enough to be moved outside. Over three days from May 31 to June 2, 2021, we removed the landscaping fabric and transferred the seedlings from the greenhouse to the treated plots. We planted 12 seedlings in each plot in a 3 × 4 array. In October of 2021 and 2022, we mowed the plots to reduce establishment of woody vegetation, which tends to outcompete herbaceous vegetation in undisturbed Eastern North American old-fields (Wright and Fridley 2010, Heckman et al. 2022).

### Plot Surveys

We surveyed the plots approximately every other week for three consecutive growing seasons. We surveyed all leaves on three haphazardly selected tillers from each plot, for an average of 9 leaves per plot. Starting with the youngest leaf, we recorded the presence or absence of herbivory damage and lesions of three focal diseases, brown patch, anthracnose, and rust. The survey was repeated for each leaf on the tiller from youngest to oldest. Older leaves are typically more damaged than younger leaves, so this survey method provides a good estimate of average damage. In the fall of each year, we identified all plant species and estimated their percent area cover within a 0.5m × 0.5m area in the center. We then estimated biomass production by harvesting all plants within a 0.25m wide strip of each plot, drying, and weighing them. In 2023, we also collected three tiller samples to test for the fungal endophyte, *Epichloë coenophiala* (Agrinostics, Phytoscreen field tiller endophyte detection kit). The seed planted was 98% positive for *E. coenophiala* and the tillers tested were 96% positive.

### 2012-2014 Experiment

To assess the generality of the effects of nutrients on disease, we also utilized data from a separate 2012-2014 experiment at Widener Farm (Heckman 2023). This experiment established 2m × 2m plots in 2012 in intact vegetation and applied factorial combinations of nutrient addition (10 g m^-2^ y^-1^ N-P-K) and combined fungicide + insecticide application, with a subplot-level leaf litter removal treatment. Early in 2013 and 2014, Heckman (2023) added seeds of 11 native species to each plot. While Heckman (2023) examined the community-level responses to these treatments and their impacts on native colonization, we sought data that was most directly comparable to the data from the 2021-2023 experiment. Therefore, we focused exclusively on data from subplots that were not sprayed with fungicide + insecticide or raked to remove leaf litter, which more closely matched the conditions in our 2021-2023 experimental plots. For the same reason, we only included the disease surveys on tall fescue leaves that were performed in 2014 in our analyses. While disease and herbivory were surveyed in 2013, this was done at the plot level rather than the sublot level. In total, we analyzed data from 20 plots in 10 spatial blocks, with one replicate of each nutrient treatment per block. In September of 2014, disease and herbivory, percent cover, and leaf nitrogen (N) concentration were quantified on tall fescue. Disease and herbivory were measured by visually estimating the percent leaf area damaged by herbivory or fungal disease on twenty haphazardly selected leaves per subplot. Percent area cover of tall fescue was estimated visually, independently of other species within subplots. Leaf nitrogen was measured by collecting three young leaves from each subplot, which were dried and pooled, then ground and quantified for leaf %N (Environmental Chemistry Lab, University of Georgia).

### Data Analysis

To analyze disease burden in the plots, we grouped surveyed leaves by plot and survey date. We then calculated the proportion of leaves surveyed with visual evidence of brown patch, anthracnose, rust, and herbivory, tracking each epidemic over each growing season (Figure S1). To integrate the prevalence of each damage type over each growing season, we calculated the area under the disease progress stairs (AUDPS, epifitter package) across the survey dates for each plot and year (Alves and Del Ponte 2021).

Given the complexity of communities, it is important to analyze the system and not just the individual relationships between plant community and disease. To assess whether the links between nutrient addition and disease were direct or indirectly mediated by some feature of the plot community, we used piecewise structural equation modeling (piecewiseSEM package) to link nutrient addition to plot level plant biomass and fescue cover and subsequently, to pathogen and herbivory damage (AUDPS) (Figure S2) (Lefcheck 2016). A direct path between nutrient addition and disease supports the nutrient-disease hypothesis, while an indirect path mediated by plot biomass or fescue abundance supports the growth-defense hypothesis. We modeled the path from anthracnose to brown patch to investigate if there were priority effects of anthracnose infection on brown patch infection as previously found in growth chamber experiments (O’Keeffe et al. 2021b, Green et al. 2024). Each equation was a linear mixed effect model with random intercepts of year and block. We log transformed biomass and percent fescue cover to improve heteroskedasticity. We also assessed correlated error between each disease and herbivory type and added the significant correlated errors to the model.

We modeled the 2014 fescue data as another, complementary piecewise structural equation model (Figure S3). We included nutrient addition as the predictor variable for leaf nitrogen, a subplot variable we did not collect in the 2021-2023 experiment, and subplot level fescue area cover. We then included models for percent leaf damage (AUDPS) of anthracnose, brown patch, rust, and herbivory with nutrient addition, leaf nitrogen, and fescue area cover as predictors. As in the 2021-2023 model, an indirect path from nutrient addition to disease that is mediated by fescue abundance supports the growth-defense tradeoff hypothesis and a direct path or an indirect path mediated by leaf nitrogen supports the nutrient-disease model. We also modeled the previously found priority effects between anthracnose to brown patch (O’Keeffe et al. 2021b, Green et al. 2024). We added plot nested within spatial block as a random intercept to each model to account for observing multiple leaves within each plot. Finally, we assessed the correlated errors between diseases and herbivory and added the significant correlations to the model.

## Results

In the 2021-2023 experiment (Figure 1A, Figure S4), soil nutrient addition directly increased plant biomass (F_2,108_ = 3.02, p = 0.003, Table S1) and brown patch disease (F_2,108_ = 2.03, p = 0.045, Table S1). Greater abundance of tall fescue increased anthracnose disease (F_2,108_ = 3.22, p = 0.001, Table S1, Figure S4). Neither fescue abundance nor anthracnose, rust, nor herbivory were directly or indirectly influenced by nutrient addition. There was no evidence of an effect of anthracnose on brown patch. There was a significant positive correlation between anthracnose and rust (F_1,108_ = 1.88, p = 0.03, Table S2). The model was a good fit to the data (Fisher’s C = 9.94, p-value = 0.446).

**Figure 1.**
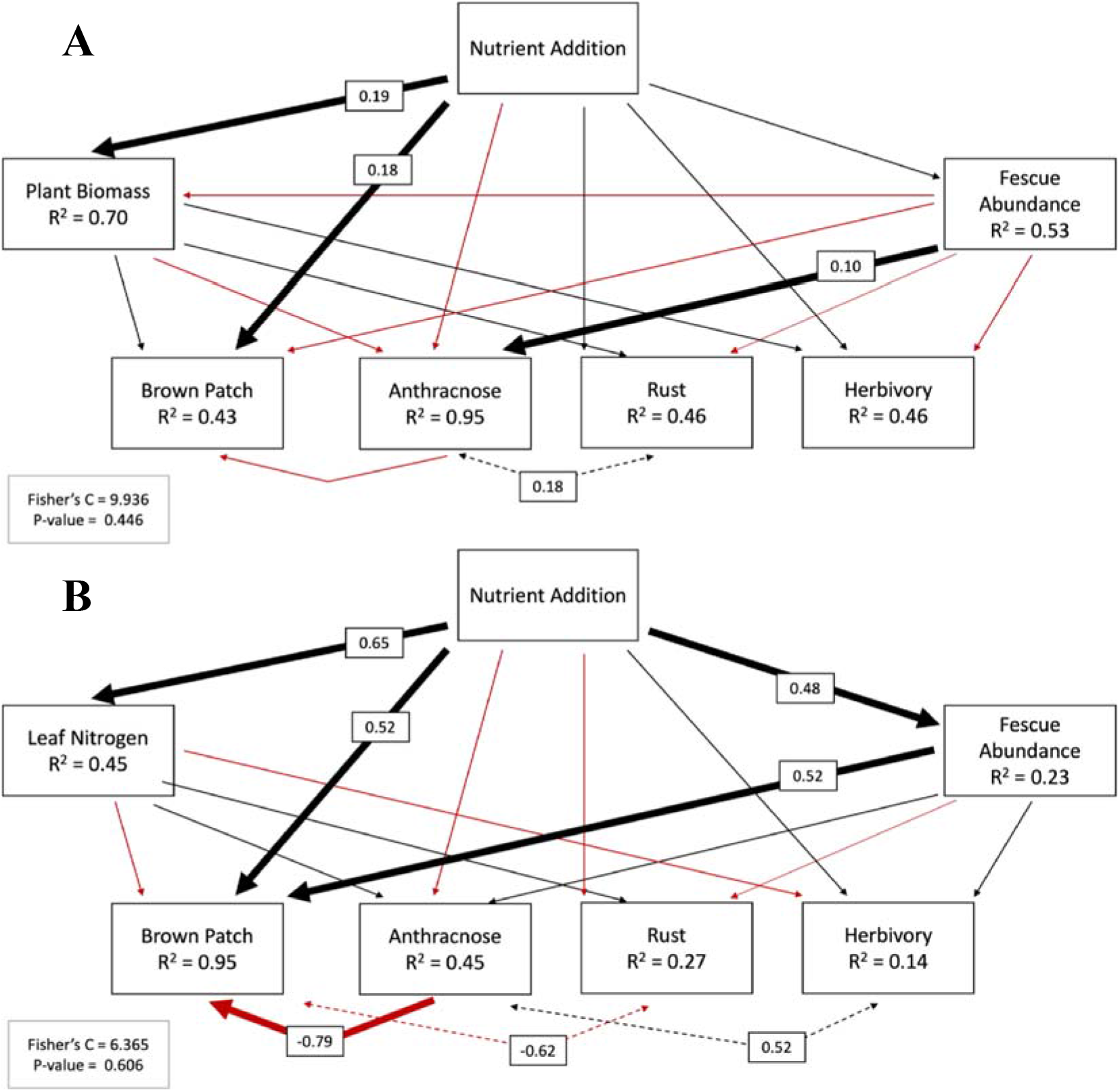
Structural equation model of the 2021-2023 (A) and 2014 (B) experiment with arrows representing paths between predictor and response variables. Red arrows represent a negative standard estimate, black a positive standard estimate and bold arrows represent a significant path. The dashed arrows represent correlated errors. Boxes on bold arrows show the significant path standard estimates. There was a significant path from nutrient addition to brown patch, partially consistent with the nutrient-disease hypothesis, and significant paths from nutrient addition to plant biomass and from fescue abundance to anthracnose burden, partially consistent with the growth-defense hypothesis. R^2^ values are conditional, accounting for the random effects of year and spatial block as well as all fixed effects.

In the 2014 experiment (Figure 1B, Figure S5), nutrient addition increased leaf nitrogen (F_1,18_ = 3.62, p = 0.002, Table S3) and fescue abundance (F_1,9_ = 2.32, p = 0.045, Table S3). Like in 2021-2023, nutrient addition directly increased brown patch disease (F_1,15_ = 2.89, p = 0.022) and it was the only disease affected by nutrient addition (Table S4, Figure S5). There was also a positive effect of tall fescue abundance on brown patch (F_1,15_ = 3.01, p = 0.016, Table S3, Figure S5), completing a positive indirect effect of nutrient addition on brown patch, mediated by fescue abundance. Anthracnose, rust, and herbivory were not affected by nutrient addition, fescue abundance, or leaf nitrogen. There was a significant negative effect of anthracnose on brown patch (F_1,15_ = −4.52, p = 0.004 Table S3, Figure S5). There were significant positive correlations between anthracnose and herbivory (F_1,20_ = 2.51, p =0.011, Table S4) and between brown patch and rust (F_1,20_ = 2.23, p =0.019, Table S4). The model was a good fit to the data (Fisher’s C = 6.365, p-value = 0.606).

## Discussion

An important challenge for ecology is to understand how effects of environmental change on infectious diseases are determined by the interplay of ecological processes at multiple levels of organization (Borer et al. 2016, Liu et al. 2017, Halliday et al. 2017, Bolnick et al. 2020).

Across biological levels of organization, our study found several intertwining effects of fertilization on the plant and pathogen communities over four years and across two separate experiments. The growth-defense hypothesis was supported by the 2014 experiment, in which nutrient addition increased host plant abundance, which in turn increased disease. The 2021-2023 experiment supported the hypothesized mechanism, but this did not explain the observed increase in disease. Similarly, nutrient addition increased leaf nitrogen in the 2014 experiment, consistent with the mechanism for the nutrient-disease hypothesis, but leaf nitrogen did not explain the observed increase in disease. Both experiments supported a direct path from nutrient addition to brown patch disease, suggesting that additional mechanisms may be important. Together, these results suggest a greater importance of mechanisms at the host population to community levels than the host individual level. Moreover, the core finding that nutrient addition increased the agriculturally and ecologically important disease brown patch was reproducible across both experiments.

Nutrient addition increased the burden of brown patch disease in both the 2014 and 2021-2023 experiments. This effect was modeled as a direct path in both experiments, and in 2014 only was joined by an indirect effect mediated by tall fescue abundance. In 2021-2023, tall fescue abundance was also positively correlated with anthracnose disease, but in that experiment nutrient addition did not increase tall fescue abundance. While not reproducible across experiments like the direct path from nutrient addition to brown patch, these effects involving tall fescue abundance provide cautious support for the growth-defense hypothesis that increasing abundance of fast-growing plants would lead to increased disease (Heckman et al. 2019, Cappelli et al. 2020).

Anthracnose burden was also correlated with increasing tall fescue abundance in our 2021-2023 experiment. The spores of anthracnose are dispersed through rain splashes from one leaf to another. Previous studies of a congener found the distance of spore dispersal to be dependent on the intensity and duration of precipitation but to generally not exceed 27cm (Ntahimpera et al. 1998). Given that spore dispersal is limited to the distance of water splashes it makes sense that we found a correlation between fescue abundance and anthracnose disease (Ntahimpera et al. 1998, Moral et al. 2012), despite the lack of effect of nutrients on abundance. Limited spore dispersal may also explain why anthracnose burden remained low in 2021, when tall fescue abundance was still low after planting earlier that year.

We failed to find support for the nutrient-disease hypothesis. While nutrient addition increased leaf nitrogen, leaf nitrogen did not increase any disease or herbivory. One explanation for the direct path between nutrient addition and brown patch is an effect of soil nutrients on soil fungi. Brown patch is caused by the fungal pathogen *Rhizoctonia solani*, which persists year-round in the soil as both hyphae and sclerotia (Ritchie et al. 2009), then in summer is transmitted from the soil to aboveground shoots. When nutrients are added to the soil, it shifts the soil microbiome allowing some species to flourish, while others are unable to compete (Deyn et al. 2004), and *R. solani* may be one of the winners under abundant nutrient conditions. Moreover, previous studies have found that nitrogen addition increased soil *R. solani* infectivity (Elmer 1997, Ritchie et al. 2009). While to our knowledge this effect has not been tested in tall fescue, an effect of nitrogen on *R. solani* abundance in soil or its infectivity may explain our result.

Finally, we did not find the facilitation of brown patch by anthracnose previously observed in growth chamber experiments (O’Keeffe et al. 2021b). In 2014, there was a significant negative effect of anthracnose on brown patch, consistent with previous experiments under field conditions (Halliday et al. 2017, Grunberg et al. 2023b). Since anthracnose arrives in the tall fescue population earlier in the growing season (Halliday et al. 2017, Grunberg et al. 2023a), this suggests a negative priority effect of anthracnose on brown patch at the level of the host population. These diverging patterns between experiments that utilized pathogen inoculations in the greenhouse and natural infections in the field suggests that there are mechanisms acting under field conditions that the growth chamber experiments do not capture. One difference is that the growth chamber experiments allowed only within-host processes, while the field experiments also allowed processes at larger biological levels, particularly transmission into and within the host population.

By integrating the frameworks of the growth-defense hypothesis and the nutrient-disease hypothesis to test the effects of soil nutrients on disease, we can draw conclusions about mechanisms at the scales of both the host community and the host individual. In the host community, our study supports that tall fescue growth is relatively sensitive to nutrient addition (Heckman et al. 2019), but that only partially explained the increases in brown patch disease under high nutrient conditions. Additionally, disease was not correlated with increased leaf nitrogen concentration within host-individuals. We thus are unable to attribute the effect of nutrients on brown patch entirely to either the growth-defense hypothesis or the nutrient-disease hypothesis. Additionally, our study highlights that disease responses to nutrients can also be shaped by interactions with other co-occurring diseases (Marchetto and Power 2018). The interactions between nutrient addition, host community assembly, and multiple pathogens is a frontier that deserves further study in order to understand and potentially mitigate the impacts of disease on wild plants and in agriculture as humans continue to alter global nutrient cycles and availability (Vitousek et al. 1997, Steffen et al. 2015).

## Acknowledgements

This work was supported by the NSF-USDA joint program in Ecology and Evolution of Infectious Diseases (USDA-NIFA AFRI grant no. 2016-67013-25762 and NSF grant DEB-2308472). Additionally, this study was funded in part by the USDA Forest Service, Rocky Mountain Research Station. The findings and conclusions of this publication are those of the authors and should not be construed to represent any official USDA or U.S. Government determination or policy. The 2014 study was funded by a Doctoral Dissertation Improvement Grant to CEM and RWH (NSF-DEB-1311289). We would like to thank I. Stiver, E. Snyder, and S. Cleary for their help with disease surveys in 2021-2023. Additionally, we would like to thank A. Hurlbert and S. McCoy for their written comments, which substantially improved this work.

