## Supplemental Material for "Amplification of disease by nutrient addition: Testing mechanisms from individual to community levels"

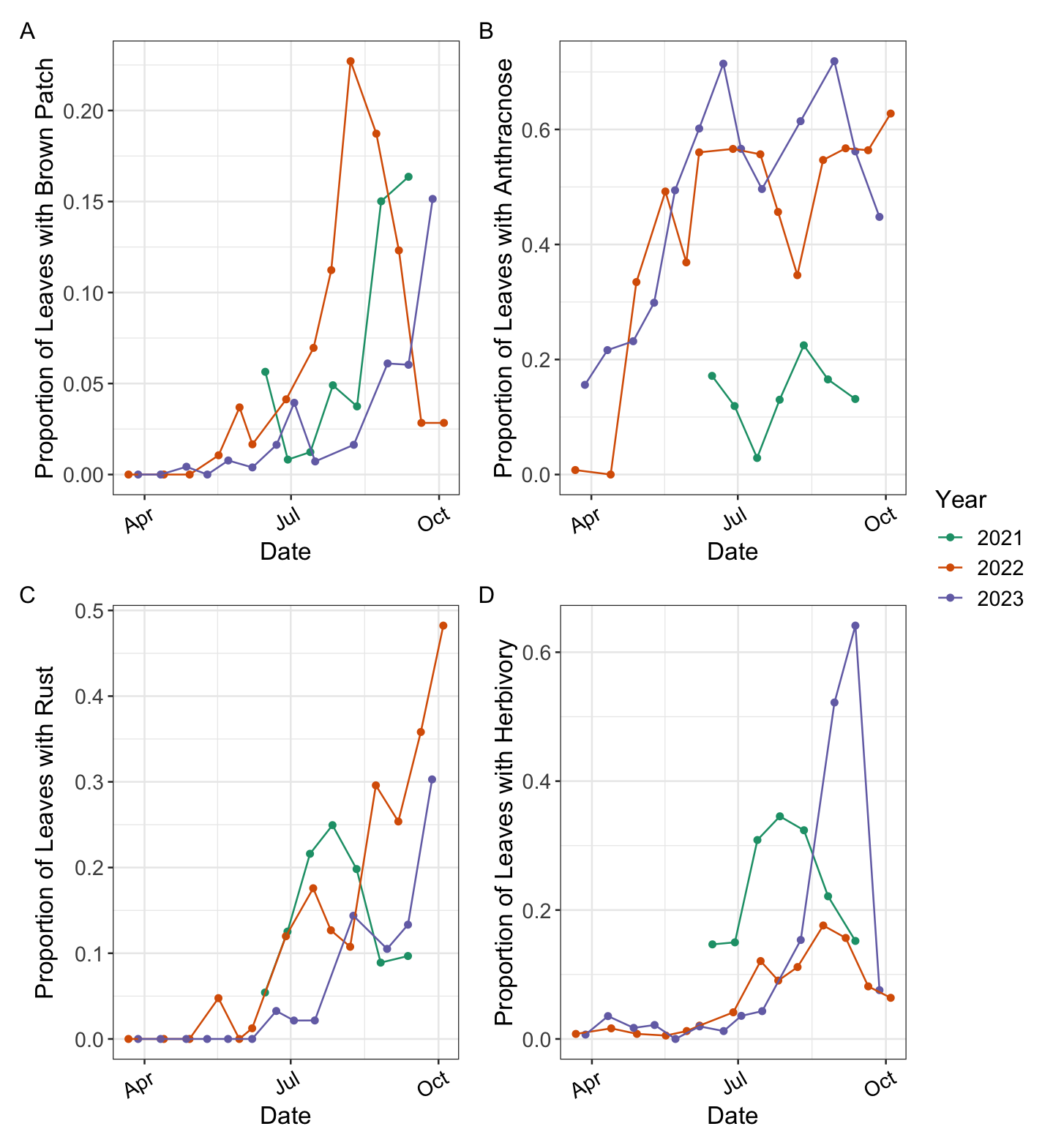

**Figure S1.** Proportion of leaves surveyed on each date with evidence of brown patch (A), anthracnose (B), rust (C), and herbivory (D) damage. The proportions are calculated across all plots within that survey date.

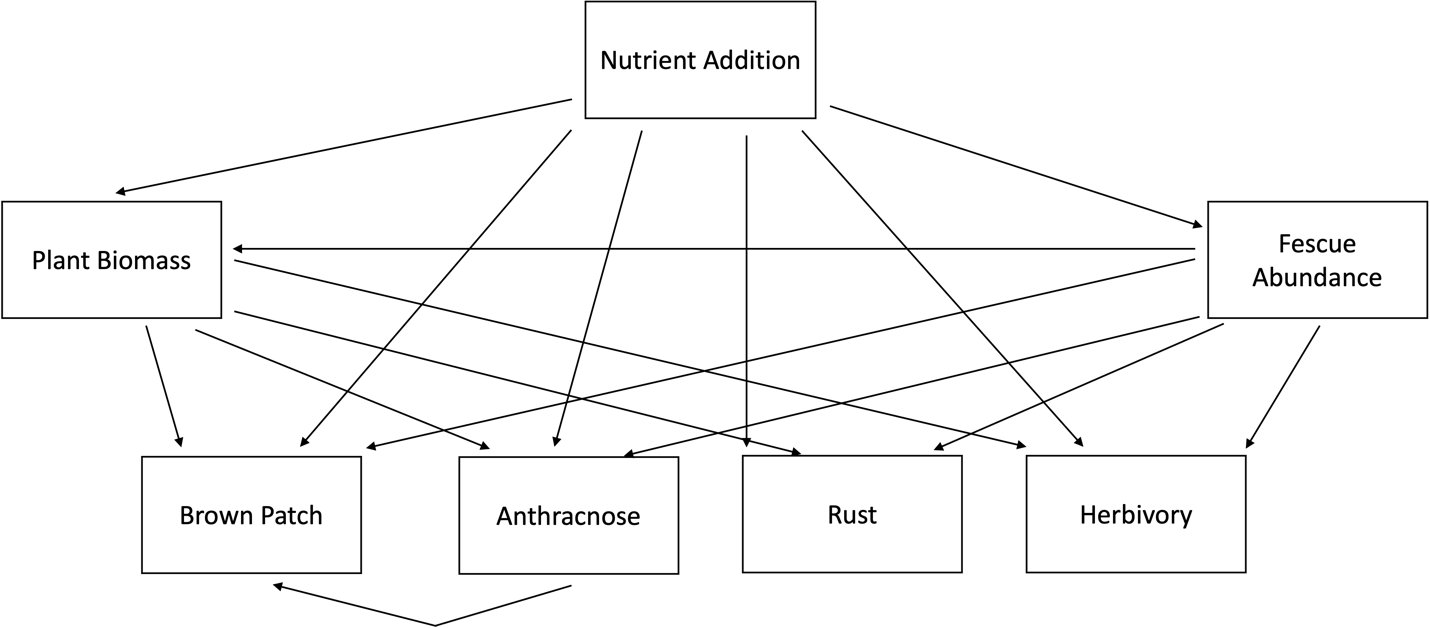

**Figure S2.** Structural equation model for 2021-2023 experiment with paths between compartments linking soil nutrients to plant biomass and fescue area cover and to disease or herbivory within the plots.

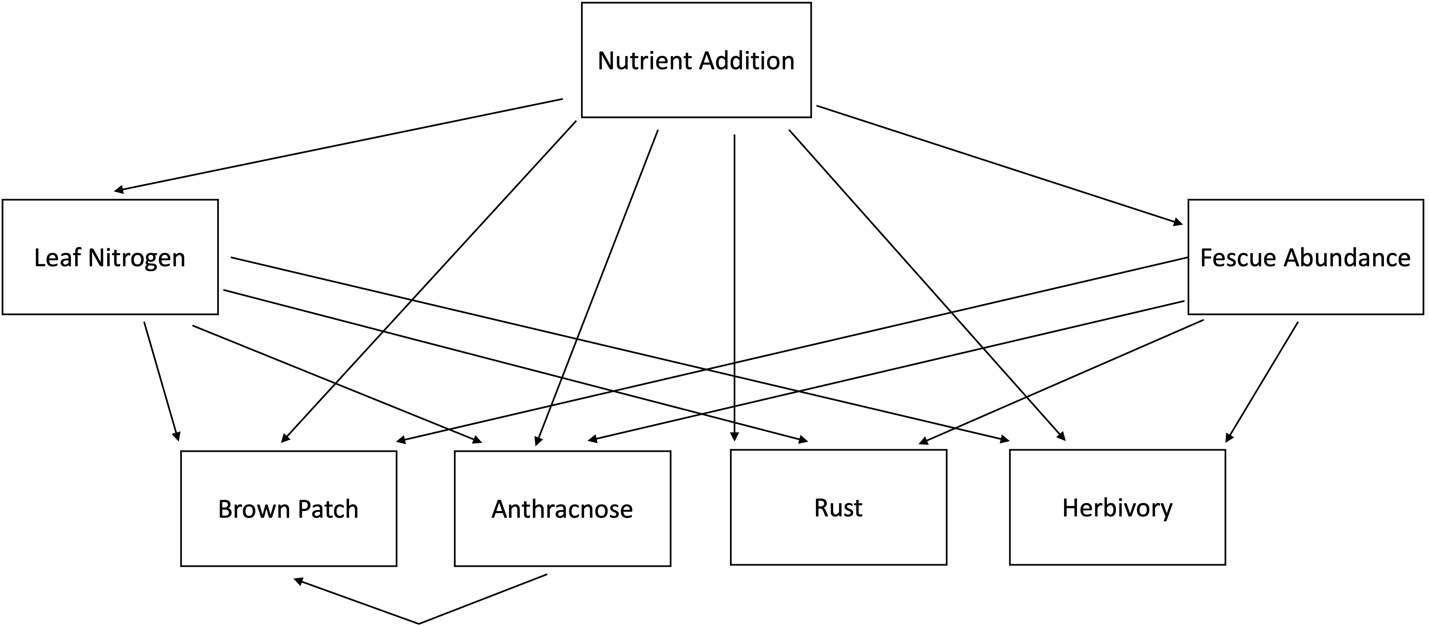

**Figure S3.** Structural equation model for 2014 experiment with paths between compartments linking soil nutrients to plant biomass and fescue area cover and to disease or herbivory within the plots.

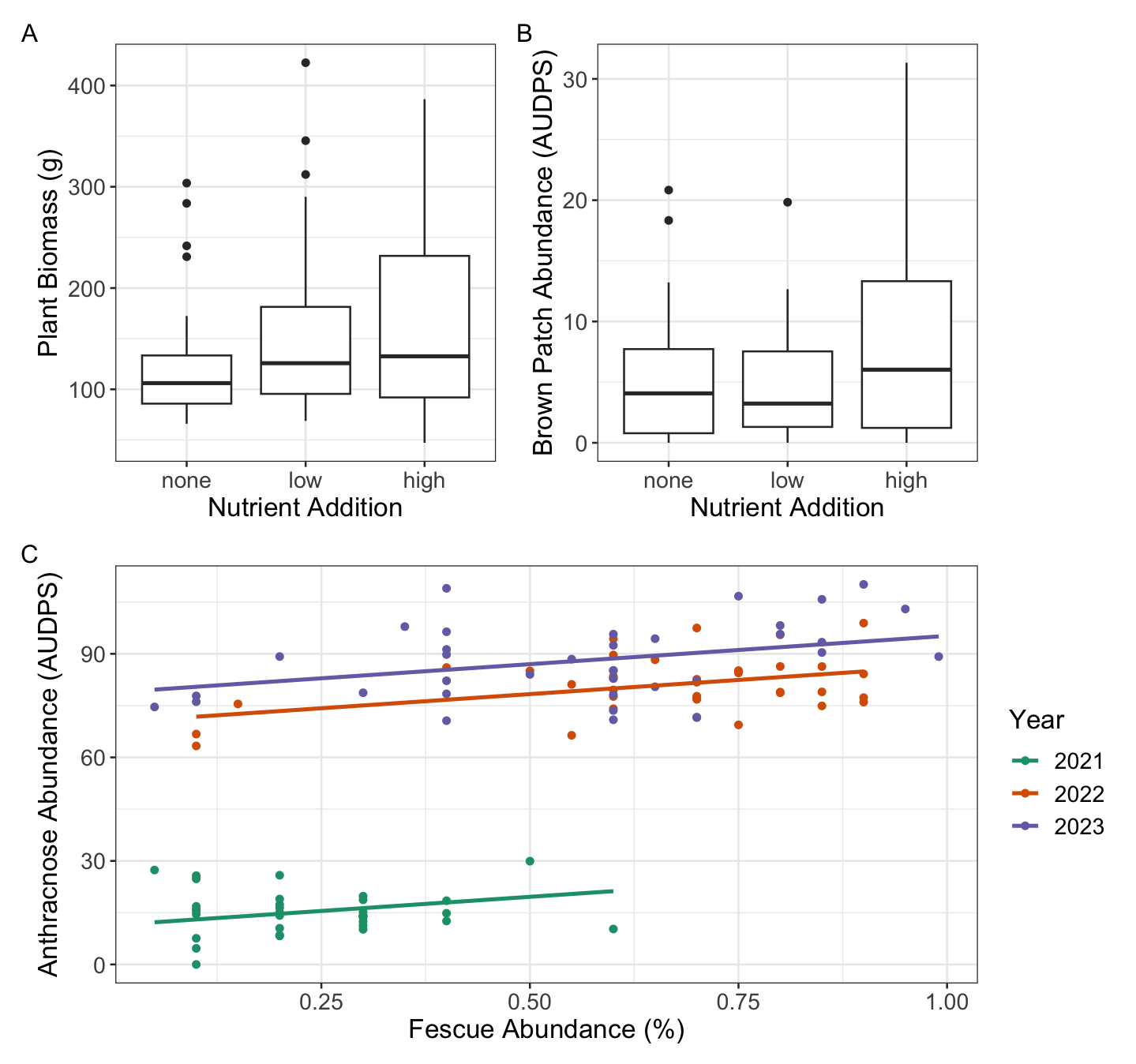

**Figure S4.** Relationships between the two variables in the significant pathways from the 2021-2023 structural equation model. Nutrient addition increased plant biomass (A) and brown patch (B), while anthracnose abundance increased with tall fescue abundance (C).

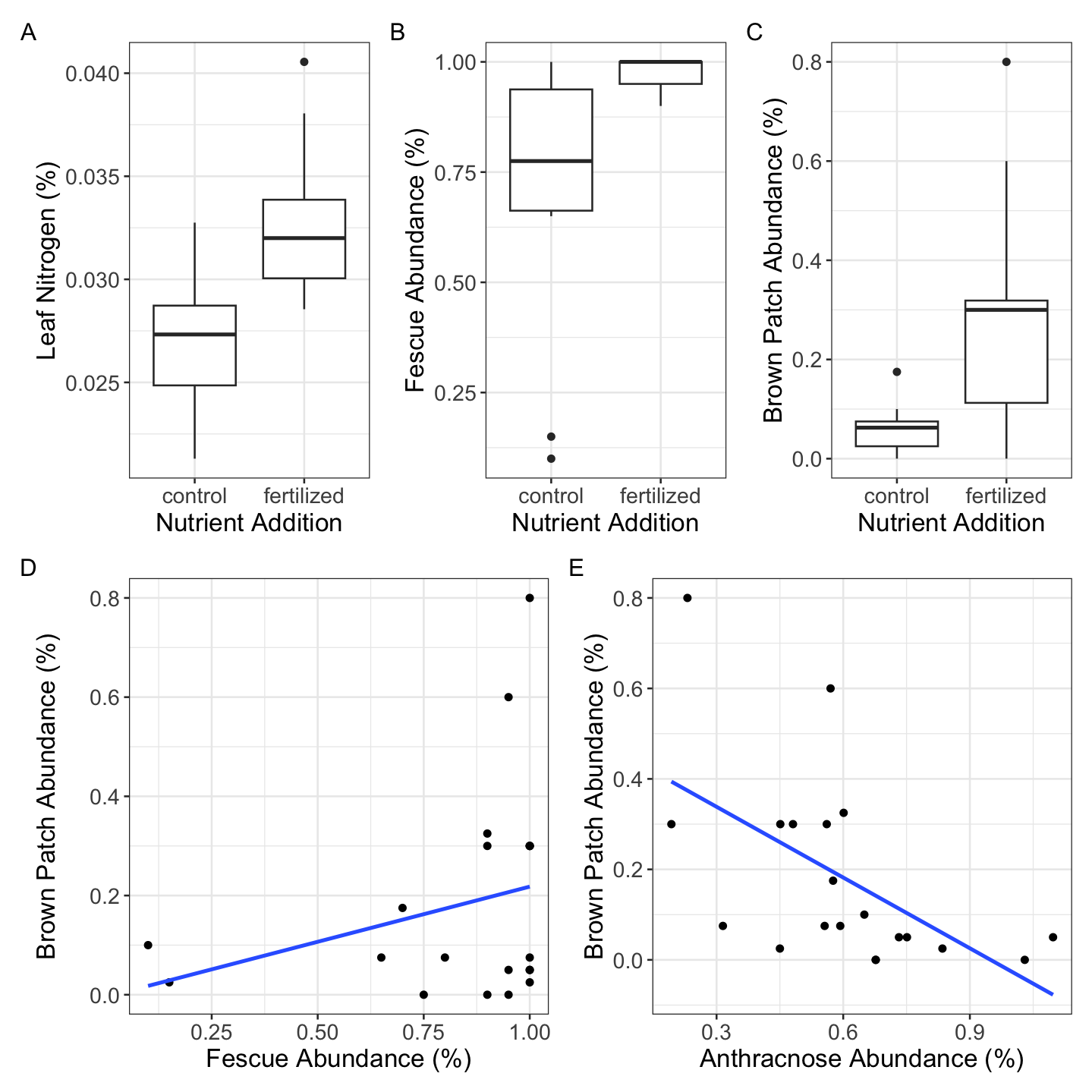

**Figure S5.** Relationships between the two variables in the significant pathways from the 2014 structural equation model. Nutrient addition increased leaf nitrogen (%) (A), tall fescue abundance (B), and brown patch abundance (C). Brown patch abundance increased with fescue abundance (D) and decreased with anthracnose abundance (E).

**Table S1.** Paths between response and predictor variables for structural equation model of 2021 – 2023 experiment (Figure 1).

| **Response** | **Predictor** | **Std Estimate** | **DF** | **F-statistic** | **p-value** |
| --- | --- | --- | --- | --- | --- |
| Biomass | Nutrients | 0.1875 | 101.009 | 3.0209 | 0.0032 |
| Biomass | Fescue Cover | -0.0644 | 102.288 | -0.8194 | 0.4145 |
| Fescue Cover | Nutrients | 0.0529 | 96.074 | 0.7267 | 0.4692 |
| Anthracnose | Nutrients | -0.0278 | 94.349 | -1.0991 | 0.2745 |
| Anthracnose | Fescue Cover | 0.1789 | 93.271 | 3.2157 | 0.0018 |
| Anthracnose | Biomass | -0.0121 | 99.591 | -0.3100 | 0.7572 |
| Brown Patch | Nutrients | 0.1789 | 95.525 | 2.0317 | 0.0450 |
| Brown Patch | Fescue Cover | -0.0719 | 87.786 | -0.6710 | 0.5040 |
| Brown Patch | Biomass | 0.2174 | 92.139 | 1.6528 | 0.1018 |
| Brown Patch | Anthracnose | -0.3975 | 22.647 | -1.3445 | 0.1921 |
| Rust | Nutrients | 0.0808 | 100.808 | 0.9610 | 0.3388 |
| Rust | Fescue Cover | -0.0327 | 101.920 | -0.3217 | 0.7483 |
| Rust | Biomass | 0.0641 | 97.350 | 0.5079 | 0.6127 |
| Herbivory | Nutrients | 0.0240 | 95.269 | 0.2995 | 0.7652 |
| Herbivory | Fescue Cover | -0.0102 | 99.593 | -0.1012 | 0.9196 |
| Herbivory | Biomass | 0.1491 | 95.292 | 1.2256 | 0.2234 |

**Table S2.** Correlated errors between pathogen and herbivory damage for structural equation model of 2021 – 2023 experiment (Figure 1).

| **Variable 1** | **Variable 2** | **Std Estimate** | **DF** | **F-statistic** | **p-value** |
| --- | --- | --- | --- | --- | --- |
| Brown Patch | Herbivory | 0.0877 | 106 | 0.8937 | 0.1867 |
| Brown Patch | Rust | 0.0871 | 106 | 0.8868 | 0.1886 |
| Anthracnose | Rust | 0.1830 | 106 | 1.8896 | 0.0308 |
| Anthracnose | Herbivory | -0.0376 | 106 | -0.3815 | 0.3518 |
| Rust | Herbivory | -0.0319 | 106 | -0.3242 | 0.3732 |

**Table S3.** Paths between response and predictor variables for structural equation model of 2014 experiment (Figure 2).

| **Response** | **Predictor** | **Std Estimate** | **DF** | **F-statistic** | **p-value** |
| --- | --- | --- | --- | --- | --- |
| %N | Nutrients | 0.6497 | 18.00 | 3.62 | 0.002 |
| Fescue Cover | Nutrients | 0.4784 | 9.00 | 2.32 | 0.045 |
| Anthracnose | Nutrients | -0.4498 | 10.25 | -1.90 | 0.085 |
| Anthracnose | Fescue Cover | 0.1807 | 15.84 | 0.88 | 0.390 |
| Anthracnose | %N | -0.4212 | 15.59 | -1.78 | 0.094 |
| Brown Patch | Nutrients | 0.5285 | 7.19 | 2.89 | 0.022 |
| Brown Patch | Fescue Cover | 0.5241 | 8.43 | 3.01 | 0.016 |
| Brown Patch | %N | 0.4486 | 8.16 | -2.27 | 0.053 |
| Brown Patch | Anthracnose | -0.7983 | 15.00 | -4.51 | 0.004 |
| Rust | Nutrients | -0.2789 | 16.00 | -0.86 | 0.401 |
| Rust | Fescue Cover | -0.2070 | 16.00 | -0.74 | 0.466 |
| Rust | %N | 0.2303 | 16.00 | 0.72 | 0.483 |
| Herbivory | Nutrients | 0.2261 | 16.00 | 0.71 | 0.486 |
| Herbivory | Fescue Cover | 0.3039 | 16.00 | 1.12 | 0.281 |
| Herbivory | %N | -0.2033 | 16.00 | -0.64 | 0.527 |

**Table S4.** Correlated errors between pathogen and herbivory damage for structural equation model of 2014 experiment (Figure 2).

| **Variable 1** | **Variable 2** | **Std Estimate** | **DF** | **F-statistic** | **p-value** |
| --- | --- | --- | --- | --- | --- |
| Anthracnose | Herbivory | 0.5207 | 20 | 2.5146 | 0.0111 |
| Anthracnose | Rust | 0.3078 | 20 | 1.3341 | 0.0999 |
| Brown Patch | Rust | -0.6277 | 20 | -3.3245 | 0.0020 |
| Rust | Herbivory | -0.0315 | 20 | -0.1299 | 0.4491 |
| Brown Patch | Herbivory | -0.3158 | 20 | -1.3724 | 0.0939 |
